# Replication is Recursion; or, Lambda: the Biological Imperative

**DOI:** 10.1101/018804

**Authors:** Scott Federhen

## Abstract

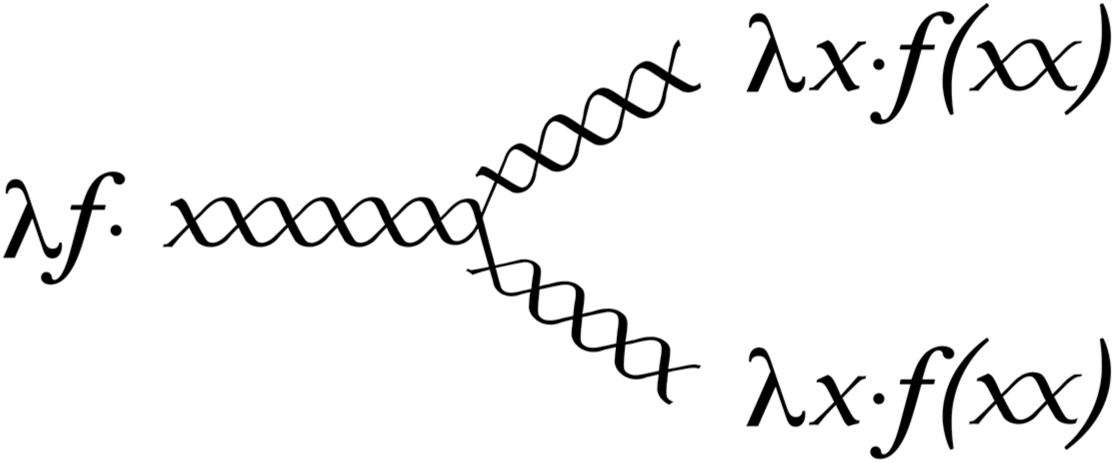

There is a striking visual symmetry between the 'paradoxical combinator' (which implements recursion in the mathematical theory of computation) and the biological replication fork (which implements reproduction in the living cell). Can this mean anything? Living organisms are information processors – the genome encodes instructions for the process of life much as computer programs encode instructions for a computer. Replication in biological systems is intuitively similar to recursion in computational systems. The following discussion will present enough of the mathematics to allow biologists to make sense of the symmetries in the logo. Mathematicians will learn nothing new about biology, but may be encouraged to look at biological processes from a new perspective.

## Computational biology: a flight of fancy

There is a fanciful role-playing game that can be played with the characters in this expression, relating to the reproduction of a living cell. λ is the lexical scoping agent in the calculus – analagous to the cell membrane, which delimits the effective domain of the genome in space. An active genome supports a function (***f***) that runs the show in a cell. This is know as ‘expressing the genome’ and can be thought of as an operating system (or executive function), implemented in the cell by machinery comprised of proteins and nucleic acids. The left-hand side of the logo shows ***f*** operating on its genome. When a cell divides, the replication fork sends a copy of the genome to each of the daughter cells, and each sets a copy of ***f*** to work on it. The analogy is not perfect – the two sub-expressions in the paradoxical combinator are not positionally equivalent; one is applied to the other. But they are absolutely interchangeable (since they are identical) so perhaps this is not disturbing enough to shatter the illusion of meaning here.

The shapes of the characters themselves display some remarkable physical properties in the light of this fancy. This is the Y-operator in the λ-calculus – the one letter from each alphabet (Roman & Greek) that actually looks like a replication fork. The standard mathematical symbol for the unknown is ***x*** – which is why it appears in the in the expression above. Self-application is at the heart of the paradoxical combinator. When the unknown ***x*** is applied to itself (***xx***) it actually looks like a unit turn of the double helix. This fortuitous resemblance allows the right-hand side of the logo to show the daughter ***f***s operating on their copies of the genome as well.

These alphabets could not have arisen by chance, but show clear evidence of intelligent character design.

## Mathematics of computation

The mathematics of computation was worked out in the 1920s & 1930s by people working on logic and the foundations of mathematics. Several different systems were developed simultaneously – recursive function theory, Turing machines, production systems, context-sensitive grammars and the lambda calculus. All of these (and any other system of sufficient power) are equivalent – they each support the same set of computable and partially computable functions.

The lambda calculus (of Alonzo Church) is in many ways the simplest and cleanest mathematical description of computation. The only object in the pure lambda calculus is the lambda expression.

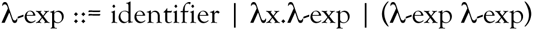

There are only three operations in the λ-calculus – application, abstraction and conversion. The workhorse is λ-application, also known as λ-reduction. A lambda expression of the form ((λx.λ-exp) λ-exp) can be ‘reduced’ by replacing all free occurrences of the identifier ‘x’ in the body of the first λ-exp with copies of the second λ-exp.

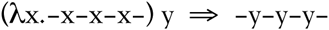

λ-abstraction is the opposite operation from λ-reduction.

λ-conversion simply allows you to change the bound variable in a λ-exp – the following to lambda expressions are equivalent.

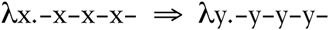

It may be necessary to do a λ-conversion in the process of λ-reduction – (λx.λy.xy) y does not reduce to λy.yy, it reduces to λz.yz (or λx.yx).

That’s absolutely all there is to the λ-calculus. A computation in the λ-calculus consists of a series of these operations on a given lambda expression, typically a series of λ-reductions. Given that the λ-calculus is powerful enough to represent all computable functions, how would one go about programming a function in the λ-calculus? There is nothing more here than a simple lexical replacement operator. How do we support recursion in the λ-calculus? The answer is a remarkable expression known as the Y-operator, the fixed-point operator, or (best of all) the ‘paradoxical combinator’. But first, a little background.

The fixed-point of a function is the set of values that do not change under the function mapping – in other words, the fixed-point of the function ***f*** is the set of all x such that ***f***(x) = x. For example, 0 & 1 are fixed-points of the function ***f***(x) = x↑2. There are no fixed-points of the function ***f***(x) = x+1, and everything is a fixed-point of the identity function ***f***(x) = x.

A fixed-point operator (**fix**) is a function that takes a function ***f*** as its argument and returns the fixed-point of that function, so that **fix**(***f***) = ***f***(**fix**(***f***)). The fixed-point operator can be defined in the λ-calculus with the deceptively simple expression:

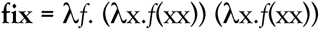

This is the fundamental operator in the mathematical theory of computation, much as replication is the fundamental operator in the biological theory of life. The appendix provides a standard treatment of logic, arithmetic and recursion in the λ-calculus, sufficient to walk through one full cycle of λ- reduction in the factorial function.

## Semantics of computation

The pure λ-calculus is endless fun for people of a certain disposition, but not everyone will want to work out division with Church numerals. Once it has been shown that it is possible to do everything with the λ-calculus, the next question becomes: is it possible to do anything useful with it?

The advent of computers and programming languages in the late 1940s and early 1950s changed the nature of computation from a mathematical abstraction to a robust and complex process in need of models. The formal syntax of programming languages was worked out in the 1950s, but it was not until the 1960s and 1970s that a formal semantics of programming languages was developed. Several different systems arose simultaneously – operational semantics akin to Turing machine models of computation, axiomatic semantics akin to production systems, and denotational semantics based on the lambda calculus model of computation together with the lattice theory as a model for data objects (Scott & Strachey). Each has its own strengths and applications, but when one uses the term “mathematical semantics” they are generally referring to the denotational model. One appealing feature of the denotational model is that it assigns meaning to arbitrary fragments of code, from the primitive elements all the way through complete programs.

*function* is a key concept in mathematics, computation and biology - it has similar but distinct meanings in each discipline. In mathematics a function is a particular kind of mapping between objects, and is subject to analysis by mathematicians. Functions in computation are related to a functions in mathematics (but tied to the idea of side-effects) and are subject to execution on a machine (a very different environment). Side-effects have no place in the primitive lambda calculus – they only appear in higher constructs that are modeled in the lambda calculus. The assignment operator is an example of a common programming language primitive that is executed primarily for its side-effect (and not for its return value) – the contents of a particular location in memory change value. This is modeled in the denotational semantics as a *store*, a function that maps locations onto values. And finally, function in biology is used to denote the jobs that proteins (and nucleic acids) do. Iteration, recursion, and looping control structures based on goto all take very similar forms when rendered as formal semantic objects in the denotational semantics, all supported by the fixed-point operator. The fixed-point operator is also sufficient to represent the complex behaviors of process forking and joining, interrupts and error processing, backtracking, and all of the exotic control structures that have appeared during the spectacular radiation of the programming languages of the the past 60 years.

The denotational semantics of programming languages is an example of a typed λ-calculus. Control structures are modelled by lambda expressions, but the data objects themselves are taken from other domains – integers, real numbers, character strings &c. From this foundation, Christopher Strachey, Dana Scott et al. erected the whole edifice of mathematical semantics (Milne & Strachey, 1975), a vast piece of mathematical engineering that makes the amusing properties of Church numerals seem like quaint pleasures from the time before computation was real work. Some programming languages are firmly grounded in the mathematics of computation – John McCarthy birthed the programming language LISP out of recursive function theory. Gerald Sussman & Guy Lewis Steele recast it (as SCHEME) in the language of the lambda calculus, in part as an attempt to implement the message-passing *actor* model of distributed computation being developed by Carl Hewitt. Other languages have semantics that are not as simple as these.

## Semantics of biological computation

Biological processes are also understood (quite explicitly) in terms of structure and function. Biological functions are mediated by molecular interactions – chemical reactions catalyzed by enzymes and transient physical binding between molecules. The genome is filled with protein binding sites – promoters & terminators, enhancers & silencers – which constitute some of the control structures that modulate the expression of the genome. Other control structures operate at levels farther removed from the genome itself. With the advent of the genomic age in bioinformatics, we might say that the syntax of biological systems is being worked out as the routine science of our day. A formal semantics for the function of structures in biological processes has not yet appeared. Functions on the genome *store* would include replacement (to model mutation, gene conversion &c.) various kinds of modifications (methylation &c.) as well as chopping and pasting (to model restriction and ligation). Primitives such as match, replace, modify, chop, ligate &c. operate on the genome; transcribe & translate create RNA & protein objects.

As an example, the prokaryotic restriction/modification system can be modeled as a set of processes that execute (**methylate** (**match** pattern)), with the side-effect that the corresponding restriction function will not cleave at that site, while other functions (e.g. transcribe & replicate) treat the modified base the same as the unmodified base. Once an organism’s genome has been protected in this fashion, it is safe to run (**cleave** (**match** pattern)) in order to terminate alien computation in the cell. Proteins that interact with the genome generally associate with the double helix in a non-site-specific fashion, and run along the molecule examining the signature of the sequence along the major and minor grooves, searching for specific patterns. This is reminiscent of pattern-directed invocation in PLANNER, although the location is the only identifier that is bound in the semantics – and reminiscent of the operation of a Turing machine.

Were we to design a programming language for this environment, these would be functions with parameters we could vary with the application. Genes code for functions in biological computation, but their products are Motie machines – each is customized to perform a specific task. EcoM1 & EcoR1 are the end products of the expression of two *E. coli* genes – *ecoM1* & *ecoR1*. Expression of these genes – (**translate** (**transcribe** *ecoM1*)) – produces a population of objects in the cell that carry out a specific instantiation of the restriction/modification system. These could be modelled as Hewitt’s *actors*, but their only possible address is their physical location in space, and they interact only with their physical neighbors (whereas the flavor of *actors* is more of computation-at-a-distance, taking at least part of its inspiration from the communication between physical particles mediated by photons & gluons &c.). Or in one of the concurrent process calculi, like the *π*-calculus (which we have rejected out of hand since *π* looks nothing like a replication fork).

The functional semantics of these protein objects can be described as:

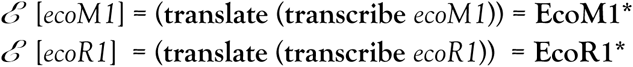

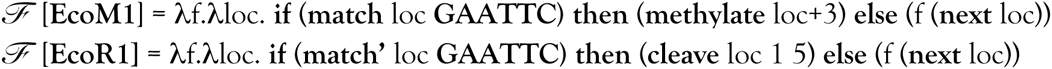

where ***ℰ*** denotes expression of the genome, ***ℱ*** denotes the function of objects in the cell, and **match’** is the pattern specific to **EcoR1**, which skips **GAA^m^TTC**. The **cleave** operation has the side-effect that (**next** loc) = **nil** for any functions operating on that locus in the genome *store*.

CRISPR is an even more complex & elegant sequence-based scheme to terminate alien computation, which has been co-opted as powerful tool for genetic engineering (as restriction enzymes were co-opted during the flowering of genetic engineering in the 1970s). The argonaute/dicer complex in the eukaryotes is a similar sequence-directed pathway that targets infecting viral sequences, but is also to regulate internal gene expression in the cell. To model these systems we need to represent the association kinetics of objects in the cell. Objects which are capable of binding each other associate and disassociate with a frequency that is determined by their relative concentrations in the cell and an association constant that is specific for that pair of objects.

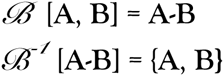

We can model the action of DNA-binding proteins the same way. As a convention, we use the ligated character ***xx*** as a dedicated variable to range only over genomic DNA. (The binding protein & genome here are reminiscent of the read head & tape of the Turing machine.)

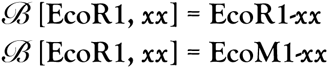

Note that we can also model enzyme-mediated chemical reactions the same way. (The cleavage of genomic DNA is of course actually an enzyme-mediated chemical reaction, though we have treated in as a purely information processing fashion.)

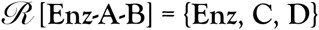

The schematic of the CRISPR locus looks as follows, where the cassettes are sequences that have been harvested from organisms whose genomes should be destroyed if they are found.

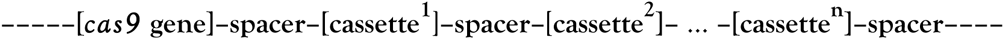

Then the semantics of the CRISPR-Cas system can be represented as:

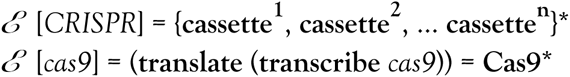

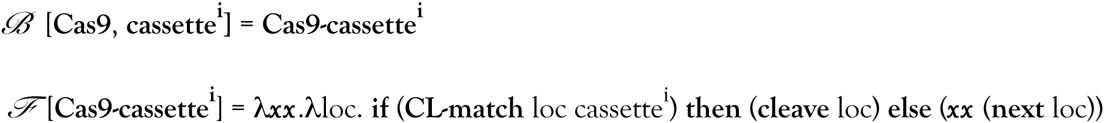

Notice that the function **CL-match** (which matches a particular CRISPR cassette with a target genome) will want to avoid recognizing the copy which is present at the host CRISPR locus (or anywhere else in the host genome – presumably these sequences are found only at the CRISPR locus in the host, but there is certain to be a perfect copy there). There is also a pathway that introduces new CL cassettes into the CRISPR locus. The more detail we know about a particular pathway, the more accurately we can describe the semantics of the functions that implement it (see Appendix III).

The genetic engineering applications of the CRISPR-Cas system were greatly advanced with the development of mutant forms of the Cas protein that only nicked the genome (rather than cleaving it) leaving it accessible for further modification.

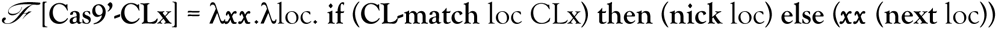

This technology has advanced to the stage where it is possible to selectively modify most locations in the genome. Recent experiments (in the yeast *Saccharomyces cerevisiae* and in the fruit fly *Drosophila melanogaster*) have shown that it is possible to construct a Cas9-primed version of a given gene that will seek out and replace the native copy on the other chromosome (analogous to gene conversion). The Cas9-primed version of the gene will then be inherited by all of the offspring, where it will convert the copy inherited from the other parent in a “mutagenic chain reaction”. This results in a hyperdominant inheritance behavior that can be used to drive a particular allele of a given gene very rapidly to uniformity in the population.

## Coda

The discrete linear structure of the genome is very compelling for a computer scientist, and much of the sequence-based information processing can be modeled in a straightforward manner. But the underlying biological processes that constitute the ‘computation’ in the life of a cell are different from programs running on machines. Any analogy can be pushed too far, and it may be that attempting to build a formal semantics of biological computation is an empty exercise akin to factoring Church numerals. But some of us have fun with that.

It may be that the binary fork is simply the simplest structure you can have, and that the duality of subject/predicate, object/action, structure/function, syntax/semantics is so pervasive in our understanding of the world that the striking symmetry between the fundamental operators of these biological & mathematical domains is not surprising, and couldn’t be otherwise.

## Acknowledgements

I was born the year after Christopher Strachey wrote some of the world’s first computer programs (for playing checkers & for playing music) on some of the world’s first computers, and the year before Watson & Crick solved the structure of DNA. These two threads have woven through my life and times like Hofstadter’s eternal golden braid. I was raised a computer scientist (Peter Bock at GW, Ed Fredkin, Gerry Sussman, Carl Hewitt & Daniel Kleitman at MIT) & on to work with the denotational semantics of AI languages with exotic control structures (Chuck Rieger, Hanan Samet & Laveen Kanal at UMd) & back to study control structures in biological systems (Bob Horvitz & Barbara Meyer, with thanks to David Botstein & Frank Solomon, again at MIT) & so on to the NCBI (Jim Ostell & David Lipman). Evidence of all of these influences is clear throughout this document. Find a way to cite Marr. And Sheree Lane, without whom nothing. Graphics by Erica Federhen.

## Appendix I – the lambda calculus

### (a) logic in the λ-calculus

In the pure λ-calculus there are neither booleans, nor integers, nor anything but λ-expressions. These are handled by providing definitions of the primitive objects and operators that behave as would be expected. These are the standard representations for boolean values in the pure λ-calculus, and some operators on them.

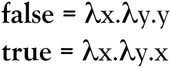

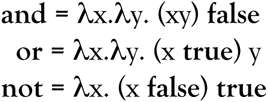

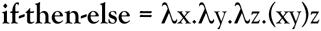

### (b) arithmetic in the λ-calculus

Studies in the foundation of mathematics tell us that all of mathematics can be reduced to arithmetic, and that arithmetic can be reduced to the number zero and the successor function, suc(x) = x+1. The representations below are known as ‘Church numerals’.

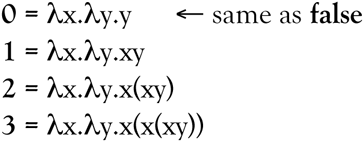

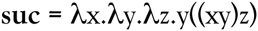

The diligent reader may want to verify that (**suc 0**) resolves to **1**.

All of the objects in the pure lambda calculus are lambda expressions, which means that they are functions that map lambda expressions onto other lambda expressions. In particular, our Church numerals are functions. What is it that they do? The number **two** takes its first argument and applies it twice to its second argument; in general, the number **n** applies its first argument to its second argument n times. We can use this insight to define a series of arithmetic functions – for example, you can add two numbers (x & y) by applying the successor function x times to the number y.

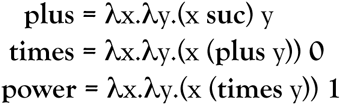

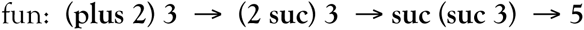

Every lambda expression can be reduced to a unique ‘normal form’ that cannot be reduced any further. Oddly enough, the normal forms for this series of arithmetic functions above get simpler and simpler.

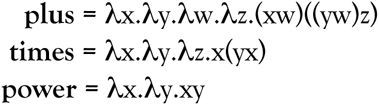

The power function (which computes y↑x) simply applies one number to the other. The function **1** is the simplest case of functional application. (When applied to two numbers, the Church numeral **1** actually computes (x↑y) – ‘unity is power’ in the λ-calculus!) Higher numbers represent higher orders of functional application, which we can write in a more familiar form:

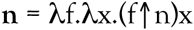

From this expression, (**n m**) → λx.(**m**↑**n**) x

This could be reduced to normal form if the appropriate Church numerals were substituted for **m** & **n**, but in its current form it is is clearly an expression that behaves like m↑n when you apply it to something. Which is all that you can ask of a lambda expression.

The successor function is defined as a primitive in the λ-calculus, Its inverse, the predecessor function, is not at all straightforward – it is easier to build λ-expressions up than it is to take them apart. The standard approach is to build the pair of numbers [n n-1] and then just select the second one when you get to the right point. Pairs of objects can be represented as:

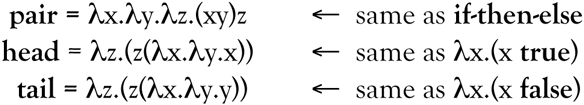

This is cons, car & cdr from LISP. There are some equivalences in the second column that may seem puzzling at first – why should the data structure **pair** have anything to do with the control structure **if-then-else**? This actually reveals something fundamental about the dual nature of lambda expressions as both data and function.

And finally, we need a definition for the function **is-zero?**, which is sufficient foundation to support a much broader equality operator for the natural numbers and beyond.

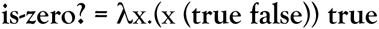

Thankfully there is no division in factorial, and the domain and range of the function is limited to the natural numbers.

### (c) recursion in the λ-calculus

The factorial function can be written as:

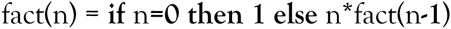

One round of λ-abstraction moves the argument to the other side of the definition:

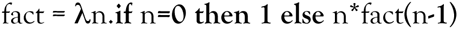

A second round of λ-abstraction gets rid of the last undefined token (fact) in the definition:

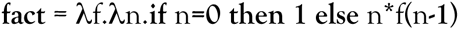

Something important has happened in this last step – the right-hand side of the ‘equation’ stands apart from the left-hand side in a way that is not true of the two preceding versions. We are left with a lambda expression that has no free variables – everything in it has been given a definition, and we are simply associating it with an arbitrary name (**fact**).

It only remains to show that (**fix fact**) **3** actually reduces to **6**.

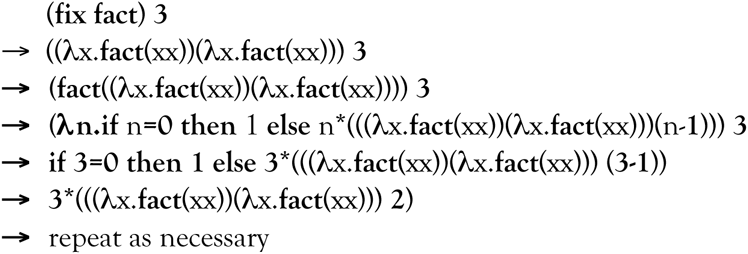

## Appendix II – a brief history of the genome

The genome and the replication operator which copies it are at the core of life. Every living cell can trace an unbroken chain of successful replications leading directly back to the origin of cellular life some 3.5 billion years ago. The binary tree structure represented by the fork in our logo is also directly reflected in the branchpoints in the evolutionary tree of life, as these also ultimately derive from the replication of the genome. In this model, currently active genomes (and the cells they run, and individuals & species that they compose) are the leading edge of a computation that has been running successfully since the origin of cellular life (and somehow on beyond that, in ways that we do not currently understand).

When you process an organism through the replication operator, you get a copy of the same thing back – in this sense, organisms are fixed points of the replication function. Except not quite – none of the extant life on earth is a copy of the primordial progenitor cell. A semantics of mutation would have to be developed for biological computation. Mutation is a side-effect, and could be modeled with an addressable genome *store* just as computer memory is modeled in the standard semantics of programming languages. There are approximately 100 de novo mutations in each haploid genome in each human generation. The same point can be made for the somatic cells in a multicellular eukaryote, when one divides you get an identical copy back – fixed points of the replication operator. Except not quite – adult organisms are not bundles of zygotes, cells differentiate as the body develops, each cell type running a different program on the same underlying genome.

There are two types of cells – haploid prokaryotic cells, and diploid eukaryotic cells, and two kinds of cell divisions – mitosis (which operates on each type of cell, and produces two identical copies of the original cell) and meiosis (a special function of eukaryotic cells which performs one replication but two rounds of cell division, producing four haploid germ cells).

Prokaryotes are simple - any two cells (from the same species or not) will trace two unbroken chains of replications back to a common progenitor. In the prokaryotes, this progression is broken by horizontal transfers of genetic data, mediated by integrative phages, conjugal plasmids, transposons &c. As a consequence, much of the evolutionary history of the prokaryotes resembles an acyclic directed graph rather than a tree (a mangrove tree of life, if you will) and the gene trees within a single species may reflect very different evolutionary histories. Genome exchange by horizontal is pervasive in the prokaryotes over evolutionary time and a number of mechanisms have evolved to support it – some bacteria actively harvest DNA sequences from the environment and incorporate them into their genomes.

In the eukaryotes, the situation is more complex. Meiosis is a special form of cell division in diploid cells, which involves a pairing between homologous (maternal & paternal) chromosomes that allows a balanced reassortment into haploid germ cells & the opportunity to recombine the product chromosomes into hybrids of the parental haploid sets. The diploid genomes of the eukaryotes support massive genomic exchange within their species envelope, and may have evolved primarily for this purpose.

Every (diploid) cell in the body can trace an unbroken chain of mitoses through the replication function back to the zygote, when the haploid maternal and paternal copies of the genome part company, and trace separate lineages back through the population. Many separate lineages actually, since each meiosis along the way fragments the genome by recombination & reassortment. The genome chunks are recognizable as ‘haplotype blocks’. The size of the genome chunks is inversely related to the distance in generations (roughly the same number of recombinations occur each generation, so the probability of recombination is proportional to sequence length). The number of genealogical ancestors increases exponentially with each generation (and would soon exceed the species population size were it not for inbreeding). One consequence is that after a small number of generations, an individual’s genetic ancestors become a vanishingly small fraction of their genealogical ancestors. After a while, the homologous fragments start to coalesce faster than they fragment, and ultimately all of the internal variation within an individual’s genome is resolved at a large number of individuals who have lived in the past. Pairs of any homologous locus that are distinguishable (differ in sequence) coalesce at individual where the mutation occurred – this is equivalent to a different ‘mitochondrial Eve’ for each locus in the genome. The average time to coalescence is proportional to the size of the mating population in the species, modified by selection, non-random mating patterns & other effects. On the average (in the human) the maternal and paternal copies of the genome will differ every thousand bases or so – regions that display a higher density of differences are separated by more evolutionary distance (i.e., they coalesce further in the past). The human genome is so well shuffled that a diploid genome from a single individual contains information sufficient to reconstruct the parameters of the population dynamics of the species for many tens of thousands of generations.

This is the purview of population genetics and the coalescent theory and is the source or yet another reason why gene trees and species trees might differ – if variation persists in the population beyond the time it takes for three (or more) species to diverge, it could settle out in ways that disagree with the corresponding species tree. If you look at the gene trees in the great apes (each of which have genomes in GenBank) the majority agree the human and chimp are sibling species, but a considerable minority support either chimp/gorilla or human/gorilla as siblings. From there, the germlines trace an unbroken chain of mitoses/meioses back through ancestral species just as in the prokaryotes, eventually passing through the great symbiosis-mediated genome merger events of the past (mitochondria & plastids being the best known examples) back to the origin of the eukaryotic cell & beyond, into the prokaryotes.

We can look forward as well as back. Once again, the prokaryotes are simple – reproduction launches a fresh copy of the genome into the world (as an independent new cell), to flourish or fade as nature sees fit. Eukaryotic cells have a different fate – for all but a handful of the germ cells (and only a small minority of those), computation will terminate with the death of the organism (if not before). With a few very peculiar exceptions – this is biology after all. The agents of transmissible venereal tumor in dogs & facial tumor disease in Tasmanian devils are examples of cancer cells that have outlived the individual they were originally a part of (a dog who lived many thousands of years ago and a devil who lived several decades ago). The component of the major histocompatiblity complex (MHC) which is responsible for rejecting tissue transplants may have evolved to combat these transmissible cancers (and not merely to frustrate 20^th^ century doctors). HeLa cells, which survive only in the laboratory, are another example of a genome that has outlived its natural span.

In contrast to the prokaryotes, eukaryotic reproduction launches a shuffled haploid copy of the two paternal genomes into the species genome pool, to fragment and scatter through the population – each locus ultimately destined either to disappear or to spread to uniformity throughout the species. Genghis Khan’s personal genome is likely to be well represented and widely distributed in today’s population – after 30+ generations the average size of these chunks should be just under two million characters (2 Mbp) by now.

To set the scale, the haploid human genome is a data store approximately 3 billion characters long, broken into 23 chromosomes. The rise between consecutive characters (base pairs) in the physical DNA molecule is 3.4 angstroms. This means that each physical copy of the human genome is about one meter long. Each of the trillions of cells in the human body has two copies of the genome. This is billions of kilometers of genome walking around with each of us, comparable to the distance from the sun to the heliopause. The genome contains 20 thousand protein coding genes, and an unknown number of additional transcribed RNAs that play a role in the expression of the genome. We are 20-30 thousand generations removed from our last common ancestor with the Neanderthal & Denisovans, but anatomically modern humans exchanged genetic material with them after leaving Africa some 2000 generations ago. We are 100 thousand generations removed from *Australopithecus*.

This brief summary has sketched the descent of extant genomes through the replication fork, but has not touched many important areas – viruses, plasmids, mobile elements within genomes, and the increasingly intimate progression from metabiomes → symbionts → endosymbionts → organelles. Many cells do bizarre things with their genomes – red blood cells throw them away, cells involved in the immune system physically rearrange them to express new protein sequences. Ciliates are single-celled, but they contain two nuclei – a germline ‘micronucleus’ which contains the genome which will be passed along to the daughter cells, and a somatic ‘macronucleus’ which contains a copy of the genome (chopped into gene-sized pieces that are circularized and linked into a mesh) which is used for expression in the genome in the cell. And we have not touched on the expression of the genome in the cell. But this is the scaffold on which a semantics of biological computation might be erected.

## Appendix III – a detailed semantics for a particular CRISPR-Cas system.

The CRISPR locus in *Streptococcus pyogenes* looks like this (figure from Chylinski et al., 2014):

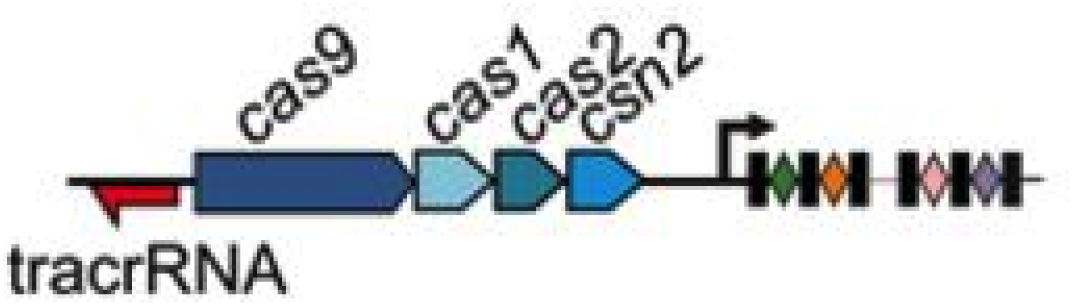

There are three protein functions (Cas9, RNase III & an unknown RNase, X) and two RNA functions (a tracer RNA and a crispr RNA) involved in implementing CRISPR immunity. The cas1, cas2 & csn2 genes encode functions involved in harvesting sequences from pathogens and adding them as cassettes to the CRISPR array. A more detailed semantics for CRISPR immunity in *Streptococcus pyogenes* follows.

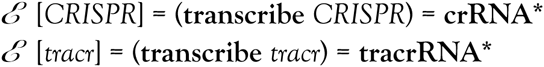

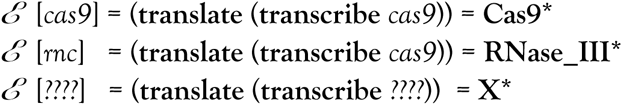

These find each other, and assemble into molecular complexes:

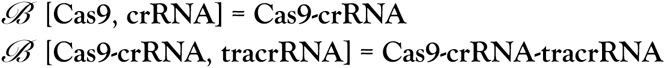

RNase III binds to this complex, cleaves the tracrRNA into a mature guide RNA, and disassociates.

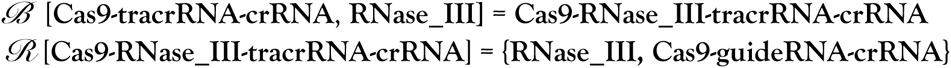

The unknown nuclease binds to the resulting complex and cleaves the crRNA to a specific cassette.

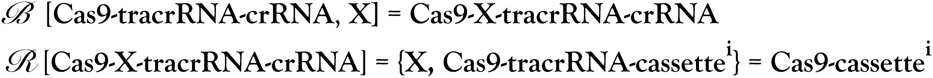

The mature enzyme complex then exhibits the same semantics as above:

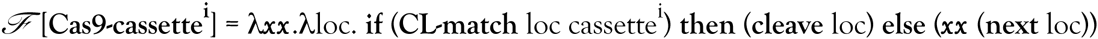

## References

Alonzo Church (1951) The Calculi of Lambda-Conversion Annals of Mathematical Studies 6, Princeton University Press.

Henk Barendregt (1984) The Lambda Calculus – Its Syntax and Semantics Studies in Logic and the Foundations of Mathematics 103, North-Holland.

Raul Rojas “A Tutorial Introduction to the Lambda Calculus” www.inf.fu-berlin.de/lehre/WS03/alpi/lambda.pdf

Robert Milne & Christopher Strachey (1975) A theory of programming language semantics [part a & b] Halsted Press.

Joseph Stoy (1977) Denotational Semantics: the Scott-Strachey Approach to Programming Language Theory MIT Press.

Scott Federhen (1981) “A Mathematical Semantics for PLANNER” Master’s Thesis, University of Maryland.

Scott Federhen (manuscript in preparation) “The Denotational Semantics of the Cell”

John McCarthy (1960) “Recursive Functions of Symbolic Expressions and Their Computation by Machine” Communications of the ACM 3(4): 184–195.

Peter Landin (1965) “Correspondence between ALGOL 60 and Church’s Lambda-notation: part I” Communications of the ACM 8(3): 89–101.

Peter Landin (1965) “Correspondence between ALGOL 60 and Church’s Lambda-notation: part II” Communications of the ACM 8(3): 158–165.

Joel Moses (1970) “The function of FUNCTION in LISP, or Why the FUNARG problem should be called the Environment Problem” SIGSAM Bulletin 15: 13–27.

Carl Hewitt, Peter Bishop & Richard Steiger (1973) “A Universal Modular ACTOR Formalism for Artificial Intelligence: IJCAI

Gerald J. Sussman & Guy Lewis Steele, Jr. (1976) “Scheme: An interpreter for Extended Lambda Calculus” MIT AI Memo 353.

Guy Lewis Steele, Jr. & Gerald Sussman (1976) “Lambda: The Ultimate Declarative” MIT AI Memo 379.

Carl Hewitt (1976) “Viewing Control Structures as Patterns of Passing Messages” MIT AI Memo 410.

Guy Lewis Steele, Fr. (1977) “Debunking the ‘Expensive Procedure Call’ Myth, or, Procedure Call Implementations Considered Harmful, or, Lambda: The Ultimate GOTO” MIT AI Memo 443.

Guy Lewis Steele, Jr. & Gerald Jay Sussman (1979) “Design of LISP-based Processors, or SCHEME: A Dielectric LISP, or Finite Memories Considered Harmful, or LAMBDA: The Ultimate Opcode” MIT AI Memo 514.

Douglas Hofstadter (1979) Gödel, Escher, Bach: An Eternal Golden Braid Basic Books.

James Watson & Francis Crick (1953) “A Structure for Dexoyribose Nucleic Acid” Nature 171(4356): 737–738.

David Marr (1969) “A theory of cerebellar cortex” J. Physiol. 202: 437–470.

Larry Niven & Jerry Pournelle (1974) The Mote in God’s Eye Simon & Schuster.

Paul Berg, David Baltimore, Sydney Brenner, Richard Roblin & Maxine Singer (1975) “Summary Statement of the Asilomar Conference on Recombinant DNA Molecules” Proc. Nat. Acad. Sci. USA 72(6): 1981–1984.

I.R. Franklin (1977) “The Distribution of the Proportion of the Genome Which Is Homozygous by Descent in Inbred Individuals” Theoretical Population Biology 11: 60–80.

Kevin Donnelly (1983) “The Probability that Related Individuals Share Some Section of Genome Identical by Descent” Theoretical Population Biology 23: 34–63.

Claudio Murgia, J.K. Pritchard, S.Y. Kim, A. Fassati & Robin A. Weiss (2006) “Clonal Origin and Evolution of a Transmissible Cancer” Cell 126(3): 477–487.

Heng Li & Richard Durbin (2011) “Inference of human population history from individual whole-genome sequence” Nature 475: 493–496.

James S. Welsh (2011) “Contagious Cancer” The Oncologist 16: 1–4.

Elizabeth A. Thompson (2013) “Identity by Descent: Variation in Meiosis, Across Genomes, and in Populations” Genetics 194: 301–326.

Rasmus Nielsen & Montgomery Slatkin (2013) An Introduction to Population Genetics: Theory and Applications Sinauer.

Svante Paabo (2014) Neanderthal Man: In Search of Lost Genomes Basic Books.

Haihua Bai et al. (2014) “The Genome of a Mongolian Individual Reveals the Genetic Imprints of Mongolians on Modern Human Populations” Genome Biol. Evol. 6(12): 3122–3136.

Stephan Schiffels & Richard Durbin (2014) “Inferring human population size and separation history from multiple genome sequences” Nature Genetics 46: 919–925

Krzysztof Chylinski, Kira Makarova, Emmanuelle Charpentier & Eugene Koonin (2014) “Classification and evolution of type II CRISPR-Cas systems” Nucleic Acids Research 42(10): 6091– 6105.

Valentino Gantz & Ethan Bier (2015) “The mutagenic chain reaction: A method for converting heterozygous to homozygous mutations” Science Express DOI: 10.1126/science.aa5945.

